# Rapid Intracellular Delivery of Human Heat Shock Protein 72 Prevents Memory Loss and Inhibits Neurodegeneration up to 9 Days After a Blast Injury

**DOI:** 10.64898/2026.06.11.726712

**Authors:** Allen Chan, Peethambaran Arun, Krutik Patel, Sarah Eintracht, Manoj Govindarajulu, Chetan Pundkar, Rex Jeya Rajkumar Samdavid Thanapaul, Gaurav Phuyal, Sonia Su, Lori Demirjian, Peyton Politewicz, Joni Ricks-Oddie, Dallas Hack, Robert Nishimura, Stephen T. Hobson, Richard A. Richieri, Courtney L. Robertson, Karolina Krasinska, Joseph B. Long, Missag H. Parseghian

**Affiliations:** Rubicon Biotechnology, Irvine, California; Blast-Induced Neurotrauma Branch, Center for Military Psychiatry and Neuroscience, Walter Reed Army Institute of Research, Silver Springs, Maryland; Center for Statistical Consulting, Dept. of Statistics, University of California, Irvine; Biostatistics, Epidemiology, and Research Design Unit, Institute for Clinical and Translational Sciences, University of California, Irvine; University of Pittsburgh, Pennsylvania; Dept. of Neurology, School of Medicine, University of California, Los Angeles; Dept. of Biology & Chemistry, Liberty University, Lynchburg, Virginia; Dept. of Anesthesiology and Critical Care Medicine, School of Medicine, Johns Hopkins University, Baltimore, Maryland; Stanford University, Palo Alto, California

**Author notes:** Dr. Chan, Dr. Arun, Mr. Patel and Ms. Eintracht, contributed equally to this work. **Name and complete address for correspondence: Missag H. Parseghian**, **PhD**, Chief Scientific Officer, Rubicon Biotechnology, 15375 Barranca Parkway, Suite B-104, Irvine, CA 92618-2205.

**Keywords:** Blast Injury, Blast Overpressure, Traumatic Brain Injury, Fv-HSP72, Heat Shock Proteins, Military Medicine, Neurotherapeutic, Neural Cytoprotection, Learning and Memory, Novel Object Recognition

## Abstract

Traumatic brain injuries (TBIs) are increasingly prevalent among military service members and are associated with long-term neurological impairment and neurodegeneration. Heat shock protein 72 (HSP72) has demonstrated cytoprotective properties and has been shown to cross the blood brain barrier in rat models of blast injury, remaining in brain tissue for up to 12 hours. In this study, we evaluate engineered Fv-HSP72 variants for their ability to reduce neurodegeneration and preserve short-term memory following blast-induced TBI. Male Sprague-Dawley rats were assigned to 9 groups of n = 8 rats. Animals were either not exposed to blast (Sham), exposed to blast (Blast Only), blast exposed and given buffer (Vehicle), or blast exposed and treated with one of three Fv-HSP72 variants, dosed at 10 or 30mg/kg at 15m post-blast. Blast exposure was generated using an Advanced Blast Simulator (ABS) producing positive static pressure to model moderate to severe blast injury. Animals were euthanized 48 hours post injury for neurodegeneration and immunologic biomarker analysis. After selecting an effective Fv-HSP72 variant using the biomarker data, additional rats were divided into Sham, Vehicle, and Fv-HSP72 treatment groups to evaluate short-term memory function through the Novel Object Recognition (NOR) test on days 2 and 8 post-blast. Analysis of cortical and spinal cord tissues demonstrated a statistically significant reduction in expression of neurodegenerative markers of Tau phosphorylation and glial injury (GFAP) for rats receiving a single dose of our clinical candidate, RBB012-CTB. In fact, the drug drove astrogliosis toward a neuroprotective state in blast exposed rats. In the NOR assay, Fv-HSP72 treated rats showed improved recognition performance, indicating preservation of short-term memory function. With similar biomarker results obtained for a controlled cortical impact injury model published elsewhere (Chan et al. *manuscript submitted*), the analyses suggest Fv-HSP72 is neuroprotective following a blast injury as well.

**One sentence summary:** This study describes the effectiveness of a biologic agent, Fv-HSP72, in significantly preventing learning and memory loss in rats for up to 9 days after a blast injury.

## Introduction

Exposure to blast overpressure and blast-induced traumatic brain injury (TBI) rose to clinical prominence during the conflicts in Iraq and Afghanistan (2001-2021) with 20% of US warfighters returning from deployment having experienced a single or multiple blast TBI^1^. The aftermath of exposure is associated with complex pathophysiological mechanisms including neuroinflammation, oxidative stress, and disruption of protein homeostasis^2^. These processes contribute to long-term neurological complications and increase the risk of neurodegenerative disorders such as dementia, Alzheimer’s and Parkinson’s diseases. For example, studies report PTSD rates in veterans lay between 11% and 20%, but they are much higher in veterans exposed to a blast TBI, with reported rates between 33% and 65%^3^. Effective management of TBI is therefore critical to mitigate both immediate neurological damage and long term cognitive and functional impairments.

Although military service members are particularly vulnerable to TBI, reporting a 3-8 fold higher incidence of neurological symptoms compared to civilian populations along with a corresponding higher prevalence and clinical burden^4^, treatment strategies remain inconsistent across acute and post-acute care settings^5^. Early clinical interventions, including timely intubation and mechanical ventilation, are often necessary to prevent hypoxia and limit secondary brain injury. However, these approaches primarily address systemic complications rather than directly targeting the underlying cellular and molecular mechanisms driving neurodegeneration.

The structural damage imposed by the nature of various forces associated with TBI release intracellular contents, such as HSPs^6,7^, into the extracellular environment and systemic circulation to trigger an immunologic response for clearing cellular debris and promoting tissue repair. These endogenous damage-associated molecular patterns (DAMPs), that activate inflammatory signaling pathways, play an important role in the cellular stress response.

Heat shock protein 72 (HSP72), once known as inducible HSP70 (HSP70i), has demonstrated potent cytoprotective activity in numerous disease and cellular stress models^8^. Studies have shown that knockdown of HSP72 using siRNA significantly impairs cellular stress tolerance^9^, highlighting its critical protective role. Following TBI, HSP72 expression is induced in the brain; however, this response may be delayed. For example, in rat models exposed to percussive head injury, HSP72 expression increases approximately 3 hours after injury in neural and glial cells at the impact site, followed by HSP72 detection at 6 hours in some blood vessels traversing the necrotic region of the impact ^10^. This delayed induction suggests that endogenous HSP72 expression may not occur rapidly enough to prevent early secondary damage following TBI.

Given the inducible and cytoprotective nature of HSP72, early exogenous delivery may provide a strategy to enhance its protective effects during the acute injury window. Conjugation with the single chain variable fragment (scFv) of monoclonal antibody 3E10, which enables targeted cellular penetration of proteins^11^, facilitates delivery of HSP72 to injured neural tissue through a compromised blood brain barrier. In this study, we investigate the neuroprotective potential of Fv-HSP72 variants and a clinical candidate’s ability to mitigate neurodegeneration and memory loss following blast-induced TBI.

## Methods

### Materials

In developing a new class of fusion proteins known as Fv-HSP72, Rubicon Biotechnology has created 6 humanized variants of the 3E10 scFv fragment and fused them to several modified forms of human HSP72 that eliminate sites of glycosylation, oxidation and acid-labile cleavage. The proteins are ∼98kD and synthesized in CHO cells, purified by chromatography, and formulated to 6mg/mL in PBS with 5% sucrose. Two humanized 3E10 variants were chosen for these studies, designated RBB010 and RBB012, based on *in vitro* DNA binding kinetics and neural cell uptake (data not shown). The 3E10 subunit was fused to the HSP72 with one of two cleavable amino acid linkers: a “Swivel” linker (RBB010, RBB012) or a cathepsin B cleavable one (RBB012-CTB). Product potency is tested both in the 3E10 subunit, which binds the target, and the HSP72 subunit, which requires a functional ATP subdomain (data not shown).

All animal studies were conducted with male Sprague-Dawley rats (CD-strain; Charles River), weighing 225-350g, at Walter Reed Army Institute of Research (WRAIR; Maryland) after review and approval by the IACUC team and US Army’s ACURO teams. Research was conducted in the AAALAC International - accredited facility at WRAIR with a Public Health Services Animal Welfare Assurance.

### Blast Injury

Male Sprague-Dawley rats were divided into 9 groups (Sham, Blast Only, Blast & Vehicle, and 10mg/kg or 30mg/kg treatment of RBB010, RBB012, RBB012-CTB) (n=8/group). The Sham control group underwent the anesthesia and handling without experiencing the blast (*i.e. uninjured & untreated*). The remaining groups of rats were exposed to two tightly coupled 20 psi blasts (two blasts within 2 min) under anesthesia in the Advanced Blast Simulator (ABS) as described earlier^12^. Drug or vehicle was injected once at 15m post-blast intravenously (IV) via tail vein injection.

### Study Groups for Screening Fv-HSP72 Variant Efficacy

Unlike the Sham group, negative control group rats were *<u>injured</u> & untreated*. There were two such groups, one receiving no injection at all (Blast Only) and another group receiving the vehicle buffer being used for Fv-HSP72 (PBS with 5% sucrose, filter sterilized). Vehicle buffer was injected intravenously (IV) 15 minutes post-blast. This second group (Blast & Vehicle) was to verify the effects of the vehicle buffer alone and each rat received a volume of buffer through the tail vein equal to the volume injected into the rats receiving drug. The remaining 6 groups of rats were *injured & treated* IV in the rat tail vein with either a 10mg/kg or 30mg/kg dose of RBB010, RBB012 or RBB012-CTB at 15m post-blast.

### Study Groups for Learning and Memory Study

Rats were randomly divided into one of three groups (Sham, Blast & Vehicle, Blast & Fv-HSP72) with 10-13 animals in each group. The Sham control group underwent the anesthesia and handling without blast exposure (uninjured & untreated). Rats were euthanized for tissue extraction on Day 2 (n=4) and Day 9 (n=6). The Blast & Vehicle rats (injured & untreated) were exposed to a blast and given vehicle buffer in the tail vein 15m post-blast at the same volume as those receiving drug. Rats were euthanized on Day 2 (n=5) or Day 9 (n=8). Those rats receiving Fv-HSP72 were injected with a 30mg/kg dose of the RBB012-CTB variant and euthanized on Day 2 (n=4) or Day 9 (n=8). All rats underwent memory testing on Day 2 with a subset of animals from each group being euthanized for biomarker analysis. The remaining rats underwent further memory testing on Day 8 and were euthanized on Day 9.

### Study Groups for Pharmacokinetic and Biodistribution Study

Fifteen “TBI” rats were put under isoflurane anesthesia and injected with Fv-HSP72 15m post-blast, while another 15 “Naïve” rats were anesthetized and received the drug without any blast exposure. Both sets of rats were divided into three groups of 5 TBI and 5 Naïve animals each. TBI and Naïve rats in Group 1 were euthanized 1h post-IV, those in Groups 2 and 3 at 4h and 12h, respectively. Brain and blood samples were extracted and shipped on dry ice to Stanford University for mass spectrometry.

### Biomarker Tissue Extraction and Analysis

Degradation of neural cytoskeletal proteins, such as α-Spectrin, have been reported to peak 48-72h post-TBI^13,14^. Although biomarkers measured in this study did not include α-Spectrin, based on these observations, our screening studies were limited to all rats being euthanized at 48h. For the learning and memory studies, a subset of rats from each group were again euthanized for tissue extraction at 48h and the remaining rats were kept for another 6 days to investigate longer-term effects of Fv-HSP72. Brain and spinal cord tissues were removed from each rat, immediately frozen in liquid nitrogen, stored at -80°C until shipment on dry ice from WRAIR to Rubicon for analysis. Tissue extraction and biomarker analysis followed protocols as described in Chan et al. (*submitted*). Tissue extracts were plated in triplicate in 96-well plates and subsequently probed with primary and secondary antibodies listed in **Table 1**. Colorimetric development of the plates with TMB substrate lasted from 5-15 minutes, as reported in each of the figure legends, before being stopped with a 1M HCl solution.

**Table 1.** List of primary (1°) and secondary (2°) antibodies used in our biomarker studies.

| Biomarker | 1°/2° | Host/ Antibody | Clone | Supplier | Catalog # | Dilution |
| --- | --- | --- | --- | --- | --- | --- |
| **3-NT** | 1° | Mouse Monoclonal | 39B6 | Sigma Millipore | #SAB5200009 | 1µg/mL |
| **S100A10** | 1° | Mouse Monoclonal | 6F4-E6-D5-C10 | Invitrogen | MA5-24769 | 1:1000 |
|  | 2° | Goat anti-Mouse Polyclonal | N/A | Jackson ImmunoResearch | 115-035-166 | 1:1000 |
| **pTau T181** | 1° | Rabbit Polyclonal | N/A | Abcam | #ab75679 | 1:2000 |
| **pTau T231** | 1° | Rabbit Monoclonal | EPR2488 | Abcam | #ab151559 | 1:5000 |
| **pTau S396** | 1° | Rabbit Monoclonal | EPR2731 | Abcam | #ab109390 | 1:2000 |
| **GFAP** | 1° | Rabbit Polyclonal | N/A | EnCor Biotechnology | RPCA-GFAP | 1:1000 |
|  | 2° | Goat anti-Rabbit Polyclonal | N/A | Jackson ImmunoResearch | 111-036-144 | 1:1000<br>1:2000 |
| **C3d** | 1° | Goat Polyclonal | N/A | R&D Systems | AF2655 | 1:40 |
|  | 2° | Donkey anti-Goat | N/A | Jackson ImmunoResearch | 705-035-147 | 1:1000 |

Statistical analysis of biomarker results used an unpaired t-test with Welch’s correction to account for any variances that are unequal between groups. Statistical significance was affirmed with p values ≤ 0.05.

### Pharmacokinetics (PK) using Liquid Chromatography (LC) - Mass Spectrometry (MS)

Brain and plasma were harvested from euthanized rats 1, 4, or 12 hrs post-injection, frozen on dry ice and shipped to Stanford for further processing. Brain tissue was weighed prior to freezing and this weight was used to normalize the mass (pg) of Fv-HSP72 protein detected per mass (mg) of brain tissue. Tissue extraction for MS, sample detection and quantitation are described in detail elsewhere [Chan et al. and Protocols.io, *both manuscript submitted*]. Statistical analysis of Fv-HSP72 uptake and its clearance over time focused on linear regression models relating peptide concentrations to time, injury group (TBI vs Naïve), and their interaction (i.e., an injury group-specific time trend). The two regression coefficients of each resulting line (y = **m**x + **b**), with **m = the slope** and **b = the y-intercept**, were used for the statistical analysis as detailed in Chan et al. (*submitted*). Linear regressions were limited to the 1h and 4h time points as most measurements of brain and blood tissue were at or below the Lower Limit of Quantitation (LLOQ) by the 12h time point.

### Novel Object Recognition (NOR) Assay

NOR tests the ability of a subject to learn and store memories in the hippocampus about a novel object^15^. The test occurred in 3 phases. In Phase 1, rats were acclimated to a chamber by allowing them to move about the box for 5 minutes the day before blast exposure. After experiencing the blast injury, each animal’s learning and memory was tested on Day 2 and Day 8. On each of those days, rats were subjected to Phase 2 where they were placed in the chamber with two identical objects and allowed 5 minutes to familiarize themselves with them. Phase 3 occurred 20 minutes later when the rats were re-introduced to the chamber; however, this time, one of the objects had been replaced with a different object of similar complexity. Over a 5-minute period, time spent investigating the novel object versus the familiar object was recorded and the Discrimination Index calculated (see text below).

## Results

### Screening Fv-HSP72 variants to identify a clinical candidate

Our central hypothesis is that the BBB is sufficiently compromised after blast exposure to permit Fv-HSP72 access to regions of tissue damage where it will protect against neurodegeneration, oxidative stress and glial cell injury. We report here three separate rat studies using the blast chamber at WRAIR that helped Rubicon Biotechnology identify a clinical candidate and demonstrate brain tissue accumulation post-blast (**Figure 1**). Nearly 20 variants of Fv-HSP72 have been engineered to improve the therapeutic’s efficacy in a microenvironment that is mildly acidic and enriched in reactive radicals^16^. Three of these variants (RBB010, RBB012, RBB012-CTB) were screened in a rat blast injury model at a 10 or 30mg/kg dose via an IV tail-vein injection 15m post-injury (**Figure 1A**). A dosage of 30mg/kg was chosen based on a tolerability study in rats that found a no adverse effect limit at the maximum achievable dose of 27.5mg/kg (data not shown). Higher dosages were achieved with improvements in the drug’s purification process; hence, 30mg/kg was tested, with a second dosage at 1/3 the amount (10mg/kg). The 3E10 antibody fragment targets extracellular DNA in regions of traumatic tissue injury and shuttles its payloads through an ENT2 channel into stressed cells; its mechanism of action having been described in multiple reports^11,17^. We tested two different fusions of the 3E10 scFv to its human HSP72 payload, both fusionsusing cleavable amino acid linkers. One of the variants (RBB012) was tested side-by-side with a “Swivel” linker (RBB012) or a proprietary “CTB” sequence cleaved by cathepsin B (RBB012-CTB), a protease that is upregulated during tissue injury^18^.

**Figure 1.**
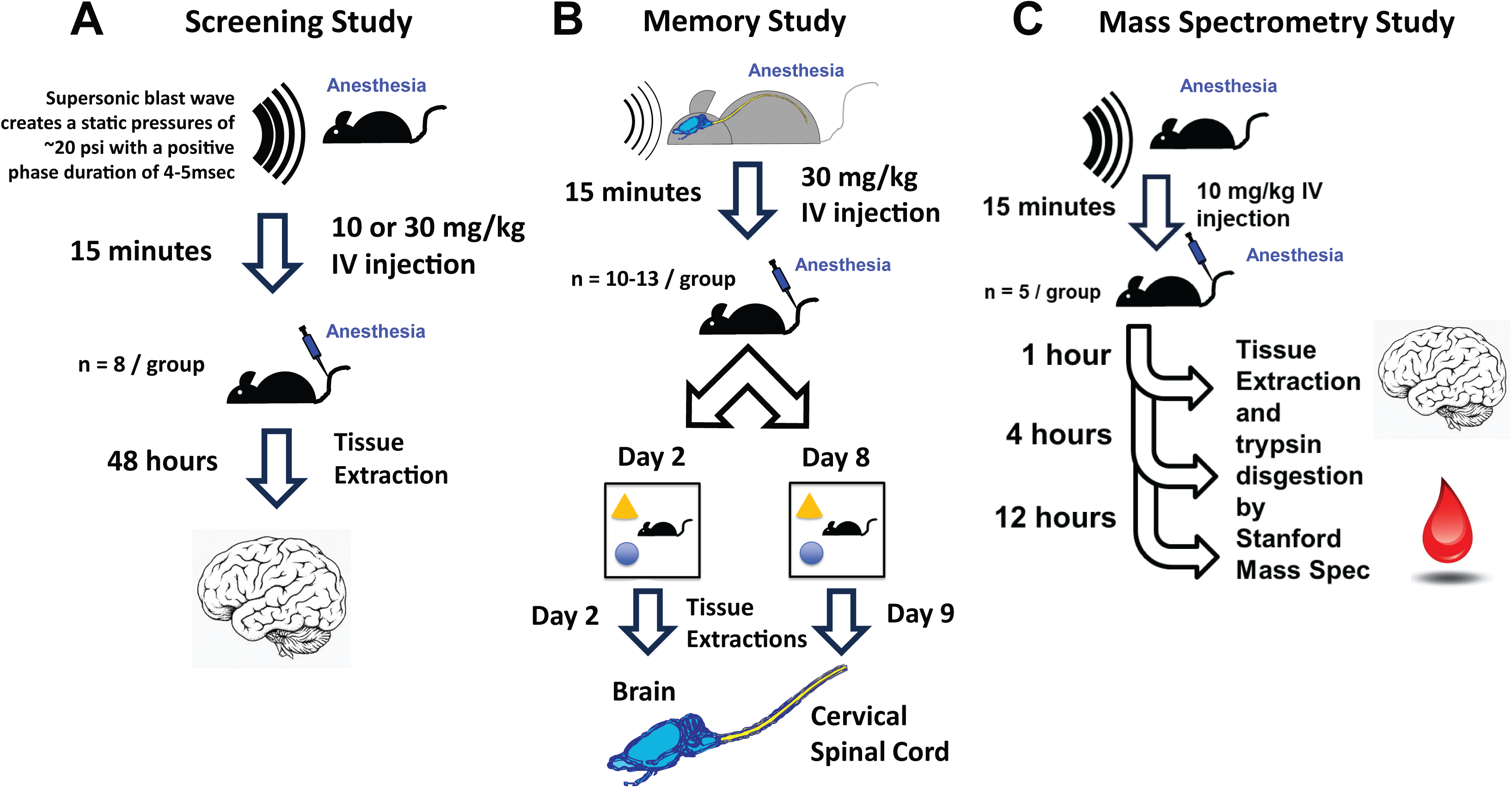
Schematics of the three separate rat studies discussed here. Anesthetized rats in each study were exposed to a blast wave of ∼20psi. **A)** Screening of multiple Fv-HSP72 primary sequence variants at two dosages (10 or 30mg/kg) are described in **Figures 2-4**. These rats were euthanized 48h post-blast, brain tissues extracted, and probed for changes in key biomarkers. **B)** Selection of a potential clinical candidate resulted in a longer study with a learning and memory test 2d and 8d post-blast (**Figure 5**). Rats only received the 30mg/kg dose and a subset of animals were euthanized on Day 2, with the remaining rats euthanized on Day 9. Brain and cervical spinal cord tissues were extracted and probed for neurodegenerative and glial biomarkers (**Figures 6-8**). **C)** A third study investigated drug uptake in the brain and blood PK using mass spectrometry. Rats received a 10mg/kg dose either 15m post-blast (TBI) or without a blast (Naïve) and were then euthanized 1, 4 or 12h post-injection (n=5/group). Brain and blood was extracted and peptides unique to Fv-HSP72 detected (**Figures 9,10**).

Blast studies conducted in the ABS chamber at WRAIR simulated a moderate to severe injury^19^ and were conducted in parallel with a team at Johns Hopkins (JHU) evaluating the same Fv-HSP72 variants to treat rats in a controlled cortical impact (CCI) model that also simulates a moderate to severe injury [Chan et al. (*submitted*)]. A kinetic focal injury to the brain can have differing mechanisms of action compared to a more diffuse injury from blast waves. This was confirmed by the different biomarkers that proved useful for screening in the two studies. Here we present biomarker examples from three parameters: oxidative stress, neurodegeneration and glial injury.

The screening studies evaluated changes in key biomarkers 48h post-blast. Secondary processes following a TBI include production of reactive oxygen and nitrogen species (ROS/ RNS). Several biomarkers are known for measuring levels of tissue damage caused by these free radicals; at 48h, measuring the nitration of tyrosine residues accumulating on proteins as 3-nitrotyrosine (**3-NT**) proved the most stable. Nitration occurs when the hydroxyl group of the tyrosine reacts with RNS, such as peroxynitrite formed from the reaction of nitric oxide and superoxide; hence, it is an indirect measure of *in vivo* radical generation. When compared to rats receiving vehicle post-blast, 3-NT levels dropped significantly for rats treated with any of the Fv-HSP72 variants at either dosage (p<0.0001, **Figure 2**).

**Figure 2.**
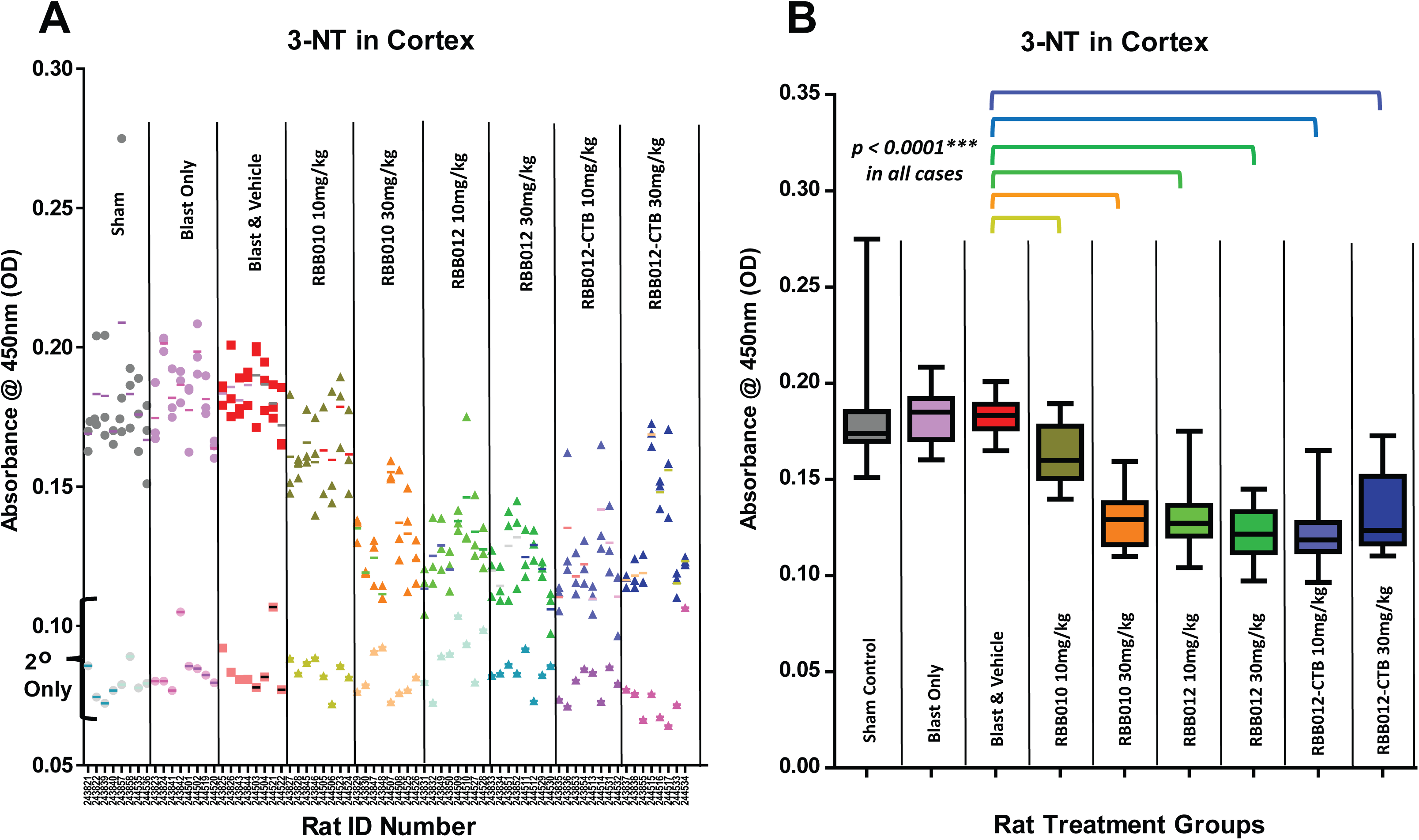
100µg of cortical tissue extract from each of the rats were plated in triplicate in the wells of an ELISA plate and probed, as described elsewhere (Chen et al. manuscript submitted), at a 1µg/mL dilution with a mouse monoclonal to the 3-NT adduct followed by a goat anti-mouse conjugated to horseradish peroxidase (GAM-HRP). TMB color development was stopped at 15m for this analysis. **A)** Scatter plot showing all of the triplicate results from each rat and their 450nm absorbance when probed for 3-NT and GAM-HRP. Scatter plots of the same rat tissue samples incubated with GAM-HRP only (2° Only) are superimposed to show baseline absorbance values. **B)** To help visualize the data easier, the same results from each group of rats are presented as box plots with statistically significant decreases in all rats treated with RBB010, RBB012, and RBB012-CTB at 10 or 30mg/kg versus the TBI & Vehicle control (p < 0.001 in all cases using a Welch’s unpaired t-test).

Our plate assay format was able to measure phosphorylation of residues on Tubulin Associated Unit (Tau) proteins, including threonine-181 and threonine-231, both markers of TBI, and serine-396, a marker strongly implicated in various tau pathologies (particularly Alzheimer’s and to a lesser extent in CTE)^20^. Given interest in developing Fv-HSP72 for clinical use in a broad series of Tau pathologies, measurement of these biomarkers illustrated a divergence in the effectiveness of the Fv-HSP72 variants with both RBB012 isoforms reducing Tau phosphorylation at S396, unlike RBB010 (compare p values in **Figure 3**). This divergence may be linked to greater cellular uptake of RBB012 versus RBB010 as measured *in vitro* with human primary neural cells. This assay involves removing any Fv-HSP72 on the extracellular surface using trypsin digestion before quantifying uptake in ethanol fixed cells using fluorescently labeled immunostains that target the 3E10 scFv subunit (data not shown).

**Figure 3.**
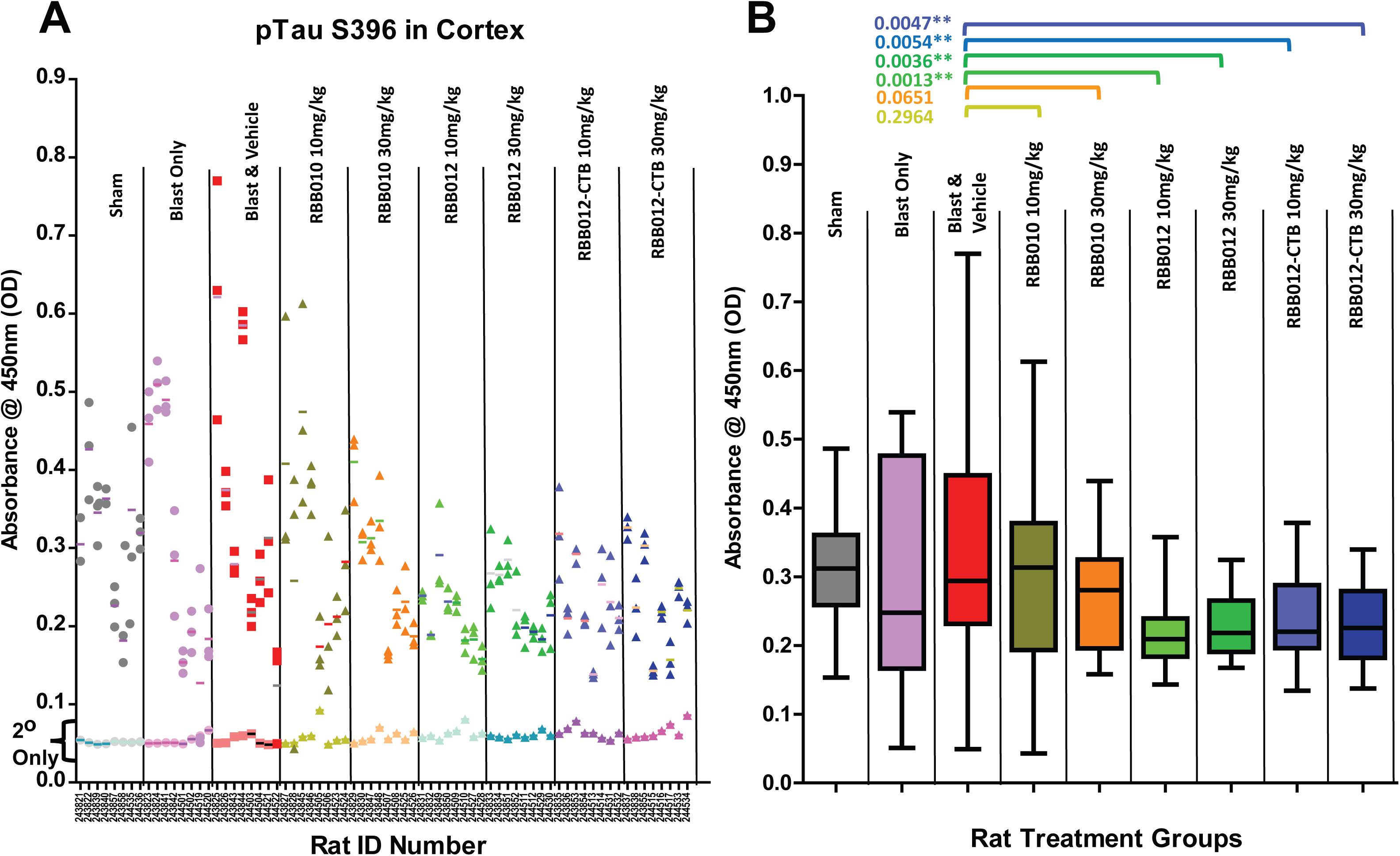
50µg of cortical tissue extract from each of the rats were plated in triplicate and probed at a 1:2000 dilution with a rabbit monoclonal to the pTau S396 phosphorylation site followed by a goat anti-rabbit conjugated to horseradish peroxidase (GAR-HRP). TMB color development was stopped at 1m for this analysis. **A)** A scatter plot showing all of the triplicate 450nm absorbances when probed for pTau S396 and GAR-HRP. When the same rat tissue samples incubated with GAR-HRP only (2° Only) are superimposed, one sees the wells probed with anti-pTau resulted in signals above background. **B)** The same data set presented as box plots show statistically significant decrease of pTau S396 in rats treated with RBB012 and RBB012-CTB of both 10mg/kg or 30mg/kg versus the TBI & Vehicle control (p < 0.05 in all cases using a Welch’s unpaired t-test).

Glial cell death is another factor leading to neurological damage post-blast injury and Glial Fibrillary Acidic Protein (GFAP) has been identified as an important marker to track the extent of cellular damage^13,21^. As was the case with measured levels of pTau S396, the Fv-HSP72 variants differed in their effectiveness in reducing GFAP levels when compared to rats receiving only a vehicle buffer (Blast & Vehicle group). Note the lack of statistical significance for RBB010 at either dose versus RBB012/RBB012-CTB at both doses (**Figure 4**). These results, along with those obtained in the parallel CCI study [Chan et al. (*submitted*)], helped identify RBB012-CTB as the highest performing candidate to advance for further development. We focused on that Fv-HSP72 variant for the remaining experiments.

**Figure 4.**
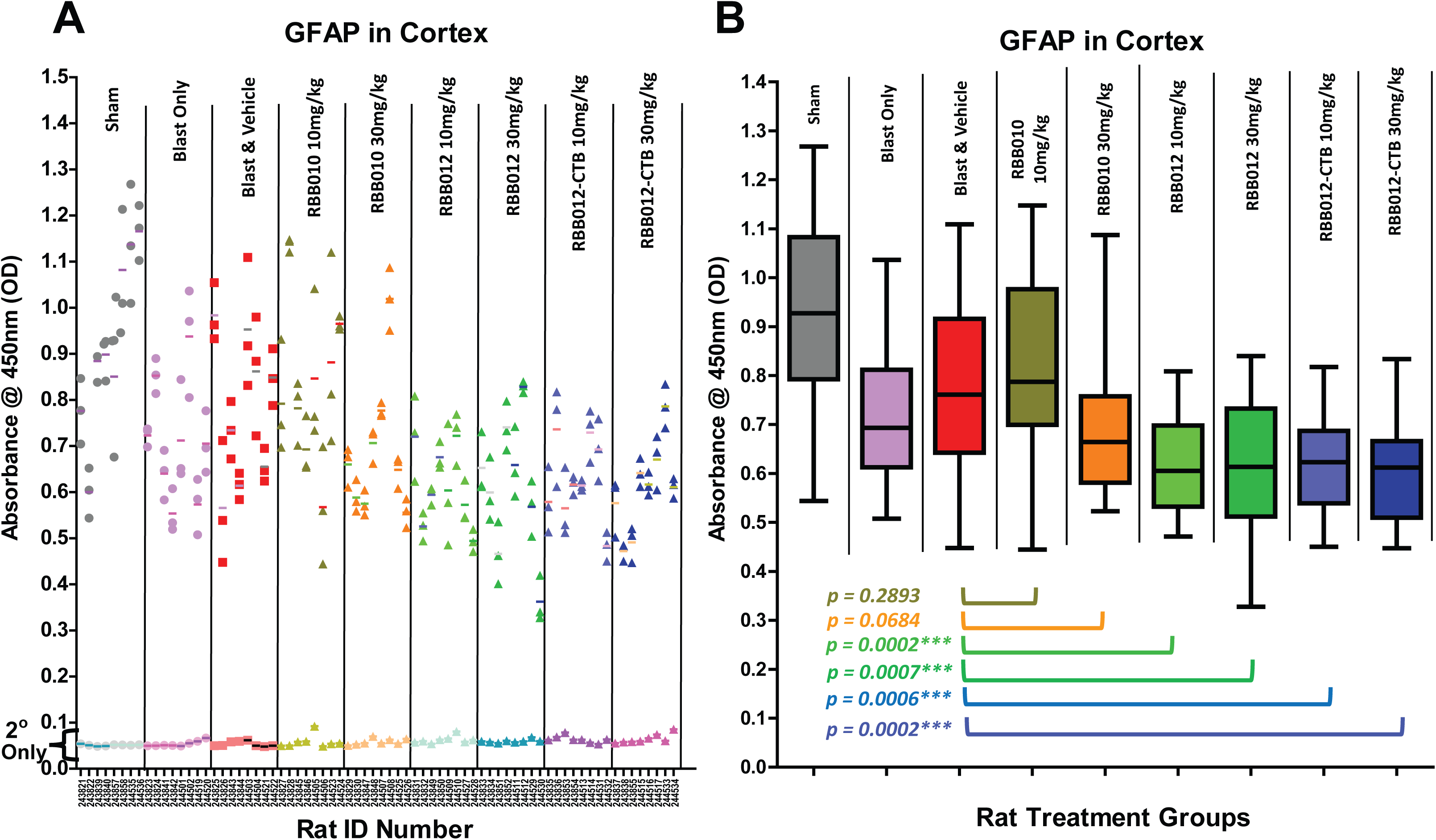
100µg of cortical tissue extract from each of the rats were plated in triplicate and probed at a 1:1000 dilution with a rabbit monoclonal to GFAP followed by a GAR-HRP secondary antibody. TMB color development was stopped at 2.5min for this analysis. **A)** A scatter plot showing all of the triplicate 450nm absorbances when probed for GFAP and GAR-HRP. When the same rat tissue samples incubated with GAR-HRP only (2° Only) are superimposed, one sees the wells probed with anti-GFAP resulted in signals above background. **B)** The same data set presented as box plots show statistically significant decreases of GFAP expression in rats treated with RBB012 and RBB012-CTB of both 10 or 30mg/kg versus the TBI & Vehicle control (p < 0.001 in all cases using a Welch’s unpaired t-test).

### Fv-HSP72 Preserves Learning and Memory after a Blast Injury

To investigate if the observed changes in TBI associated biomarkers result in corresponding behavioral impact, a new set of rats were exposed to the same blast parameters in the ABS chamber and then subjected to NOR tests to evaluate learning and memory storage in the hippocampus (**Figure 1B**). The test occurs in 3 phases; Phase I: chamber acclimation; Phase 2: object familiarization; Phase 3: evaluation of novel objects. In Phase 2, rats familiarize themselves with two identical objects in a chamber for 5 minutes before being returned to their cages for 20 minutes. During this time, one of the objects is replaced with a novel one, similar in complexity (**Figure 5A**). Once they are returned to the chamber in Phase 3, rats who have already familiarized themselves with the original object are expected to spend most of the time learning about the novel object. Rats that are memory impaired may spend an equal or greater amount of time refamiliarizing themselves with the original object. Time measurements are normalized in the form of a Discrimination Index, where the difference in time spent with a novel object and a familiar object is divided by the total time exploring both objects.

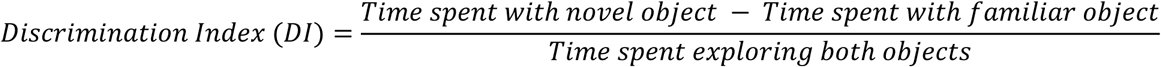

**Figure 5.**
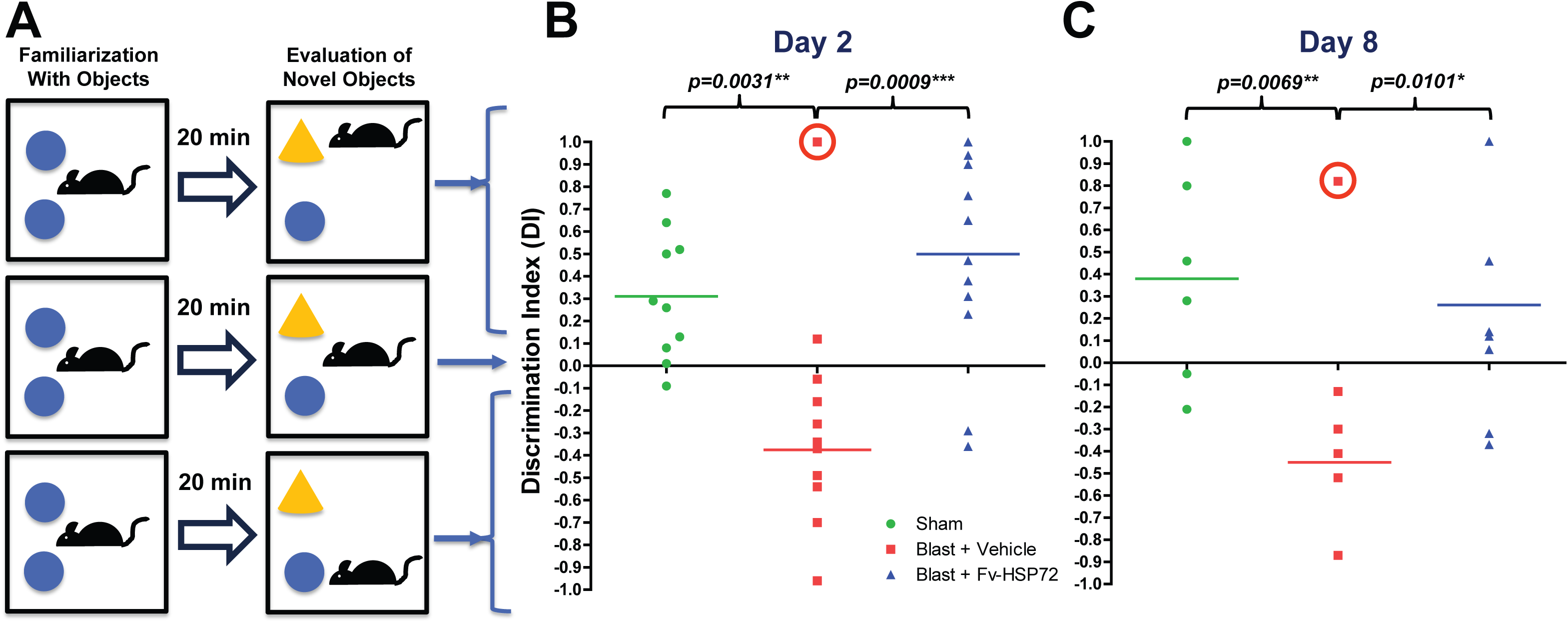
NOR Results. **A)** Schematic of Phases 2 (Familiarization) and 3 (Evaluation) of the NOR task helps interpret the Discrimination Index. Values equaling 0 represent no preference for the familiar object or the novel one. Positive values indicate preference for the novel object, while negative values indicate preference for the familiar one. **B)** Discrimination index values measured 2d post-injury are shown as a scatter plot. Results were statistically significant regardless if the one outlier from the Blast + Vehicle Control group, which was 270% of the mean, was included or not (circled). Blast injured rats only receiving the vehicle buffer spent most of their time refamiliarizing themselves with the original object they had been introduced to in Phase 2. When compared to the Vehicle group in a Welch’s unpaired t-test, most of the Sham (p=0.0031) and Blast + Fv-HSP72 treated (p=0.0009) rats focused on the novel object. Removal of the outlier resulted in p<0.0001 for both the Sham and Blast + Fv-HSP72 groups. A one-way ANOVA also was highly significant with (p=0.0014) or without (p=0.0002) the outlier. **C)** 8d post-blast, rats treated with Fv-HSP72 still showed a greater DI score than their Blast + Vehicle counterparts with p=0.0101 after a 238% outlier from the group mean (circled) was dropped. One-way ANOVA of the Day 8 data was also significant (p=0.0188). The mean for each group is represented by a line. The mean for the Blast + Vehicle reflect the data without the outliers. * p ≤ 0.05, ** p ≤ 0.01, *** p ≤ 0.001.

A single injection of RBB012-CTB at 30mg/kg 15m post-blast injury resulted in rats retaining their ability to remember which objects are familiar to them and which are novel (**Figure 5B**). The ability to retain memories on par with sham control rats lasted for up to 8d after the blast injury (**Figure 5C**). There was no statistical difference in the performance of the Sham group and the Blast + Fv-HSP72 rats on either day.

As stated in Methods, a subset of rats from all 3 groups were euthanized on Day 2 and the remainder on Day 9 for tissue extractions from the brain and the cervical spinal cord (**Figure 1B**). ELISA tissue analysis allowed us to examine if biomarker levels at 48 hours post-blast were still relevant 9 days later; thus, providing a potential explanation for the improved performance of rats treated with RBB012-CTB versus vehicle only. Tau phosphorylation biomarkers considered important in neurodegenerative pathways due to TBI (T181, T231) and Alzheimer’s (S396) were significantly decreased in the cortex (**Figure 6**) and, to a lesser extent, the spinal cord (**Figure 7**) in those rats receiving Fv-HSP72. However, results were not identical for the two parts of the central nervous system possibly indicating differences in the distribution of damage caused by the head-on orientation of the rats when exposed to the blast; thus, resulting in a differential distribution of our drug in areas with greater injury.

**Figure 6.**
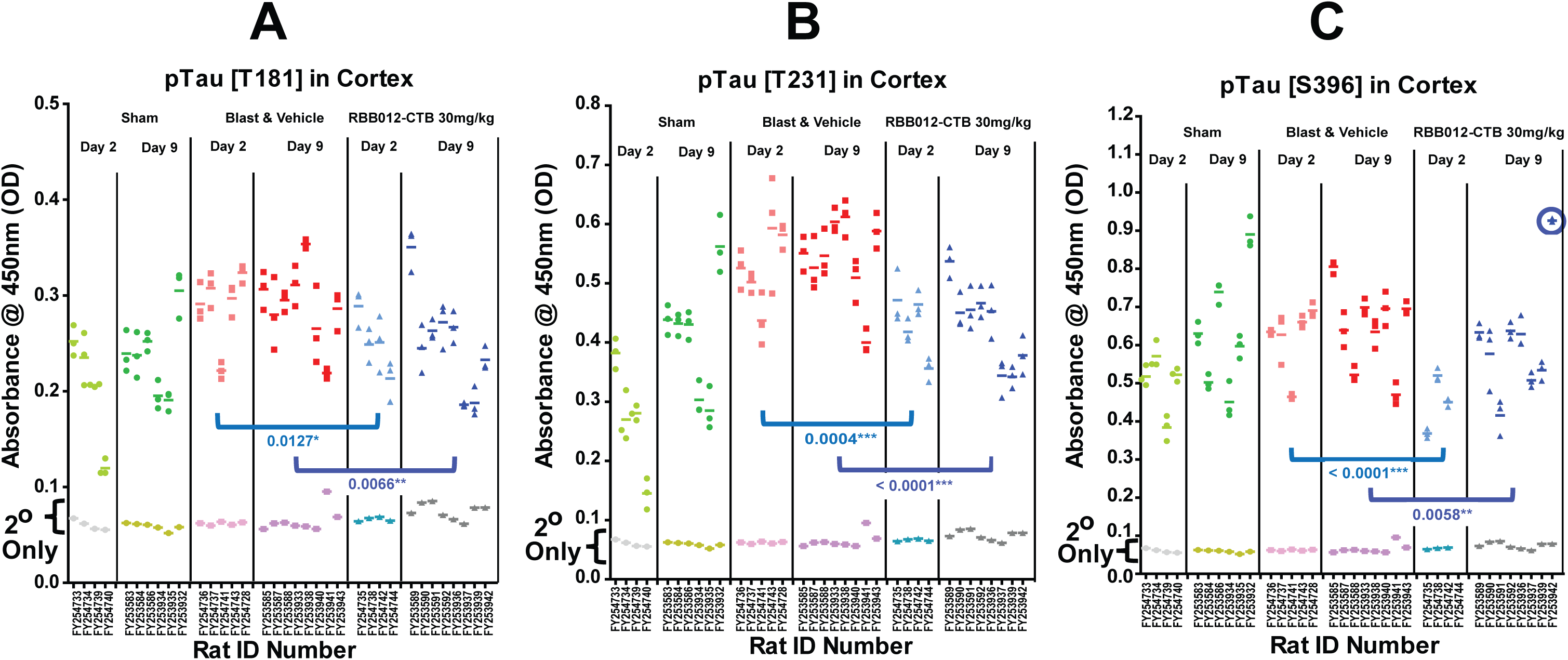
50µg of cortical tissue extract from each of the rats were plated in triplicate in the wells of an ELISA plate and probed with rabbit antibodies (**Table 1**) to phosphorylated Tau **A)** at Threonine-181 (T181, 1:2000 dilution); **B)** at Threonine-231 (T231, 1:5000); and **C)** at Serine-396 (S396, 1:2000). All three primary antibodies were detected with a GAR-HRP secondary at a 1:1000 dilution. TMB color development was stopped at **A)** 2m for T181; **B)** 1m for T231; and **C)** 1m for S396. Scatter plots of the 450nm absorbances summarize the results from each rat in the Sham (green), TBI + Vehicle (red) and RBB012-CTB treated (blue) groups. Biomarker results from tissues harvested on Day 2 are presented in lighter colors and their Day 9 counterparts are presented in the corresponding bold colors. The same rat tissue samples incubated with GAR-HRP only (2° Only) are superimposed. Mean absorbance values for rats receiving Fv-HSP72 treatment at 15m post-blast are significantly lower on Days 2 and 9 compared to rats receiving vehicle only, across the board, for all three neurodegeneration biomarkers. One outlier was dropped in the RBB012-CTB treated rats that was 153% from the group mean (circled).

**Figure 7.**
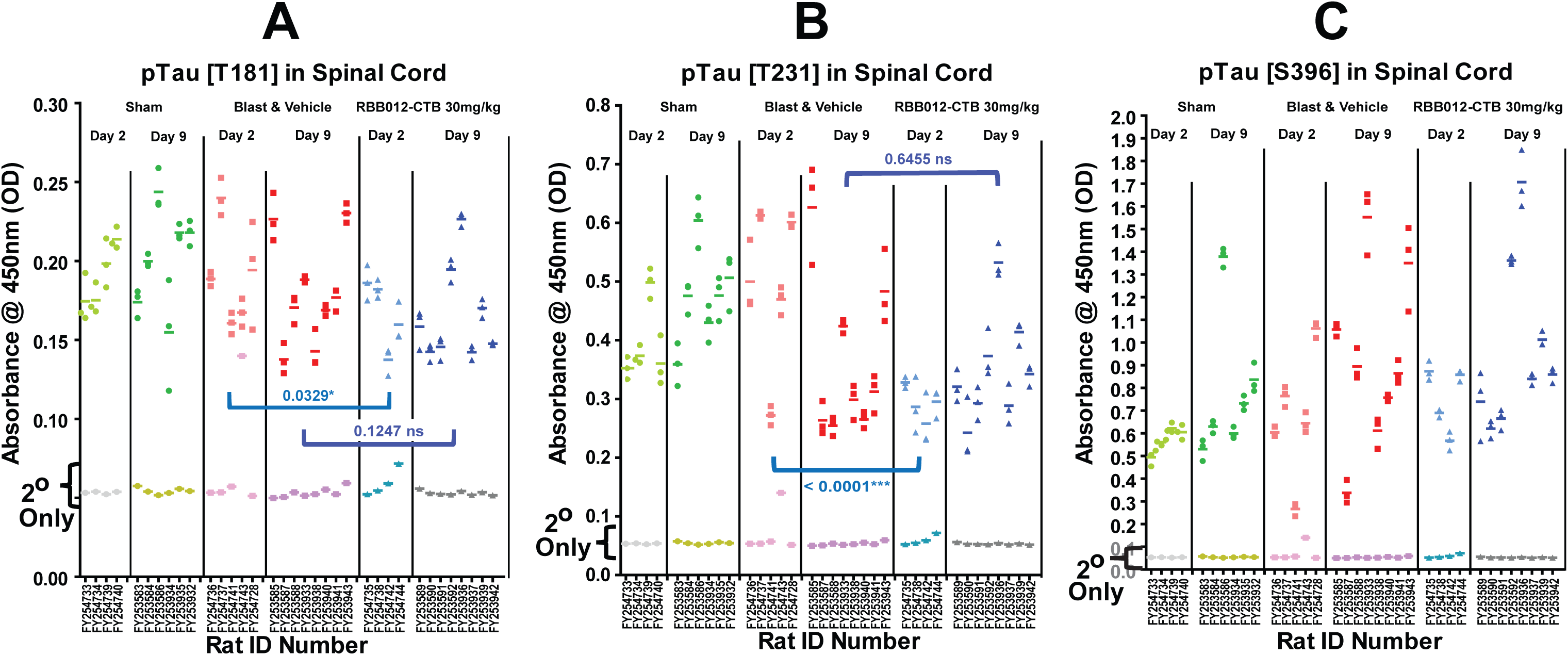
50µg of spinal cord tissue extract from each of the rats were plated in triplicate in the wells of an ELISA plate and probed with the same antibodies and dlutions as in **Figure 6**. TMB color development was stopped at **A)** 10m for T181; **B)** 3m for T231; and **C)** 4m for S396. Statistically significant decreases in pTau signals for Fv-HSP72 treated versus untreated rats were only seen on Day 2 for the TBI markers **A)** T181 and **B)** T231. The differences disappear by Day 9 due to a decrease in the mean phosphorylation levels from rats treated with vehicle control, rather than an increase in levels for those rats treated with RBB012-CTB. **C)** The Alzheimer/CTE biomarker (S396) did not show any significant difference between Fv-HSP72 treated and vehicle control rats in the spinal cord. ns = not significant.

Glial cell injury and astrogliosis occurring in the first few hours and days after a TBI are considered important contributors to neurological damage^13^. The level of glial injury was measured using an antibody targeting GFAP (**Figure 8A,B**) and the activation state of the most abundant group of glial cells, the astrocytes, was elucidated (**Figure 8C-F**). As was the case with the 48h GFAP results in our screening study (**Figure 4**), a single 30mg/kg dose of RBB012-CTB significantly reduced GFAP levels in the cortex at 2d, and up to 9d, post-blast when compared to rats receiving vehicle only (**Figure 8B**, p<0.0001). Astrocytes have pleiotropic roles supporting neuronal function in the brain and, upon injury, can undergo astrogliosis – the activation into either a neuroprotective “A2” state that promotes tissue repair or a neurotoxic “A1” state that promotes inflammation. The A2 state is detected by probing for the S100A10 calcium binding protein (**Figure 8C,D**). RBB012-CTB given to rats 15m post-blast resulted in significant activation of neuroprotective astrocytes for up to 9d when compared to a vehicle control (p<0.0001). The A1 state is detected by probing for the C3d complement factor (**Figure 8E,F**). Statistically significant increases are seen of this neurotoxic marker, although the increase may not be biologically meaningful given the slight increase in absorbance values above baseline for both the vehicle control and the Fv-HSP72 treated animals after 15m of TMB color development. The average absorbances obtained for C3d were only 1.2-fold and 1.5-fold greater than the average baseline absorbances (2° Only) for Blast & Vehicle and RBB012-CTB treated rats, respectively. As a comparison, average absorbance signals obtained for S100A10 with 8m of TMB development were 5.5-fold and 7.3-fold greater than baseline for Blast & Vehicle and RBB012-CTB treated rats, respectively (**Figure 8C**). Thus, it appears astrogliosis largely favored the neuroprotective A2 state in those rats given RBB012-CTB.

**Figure 8.**
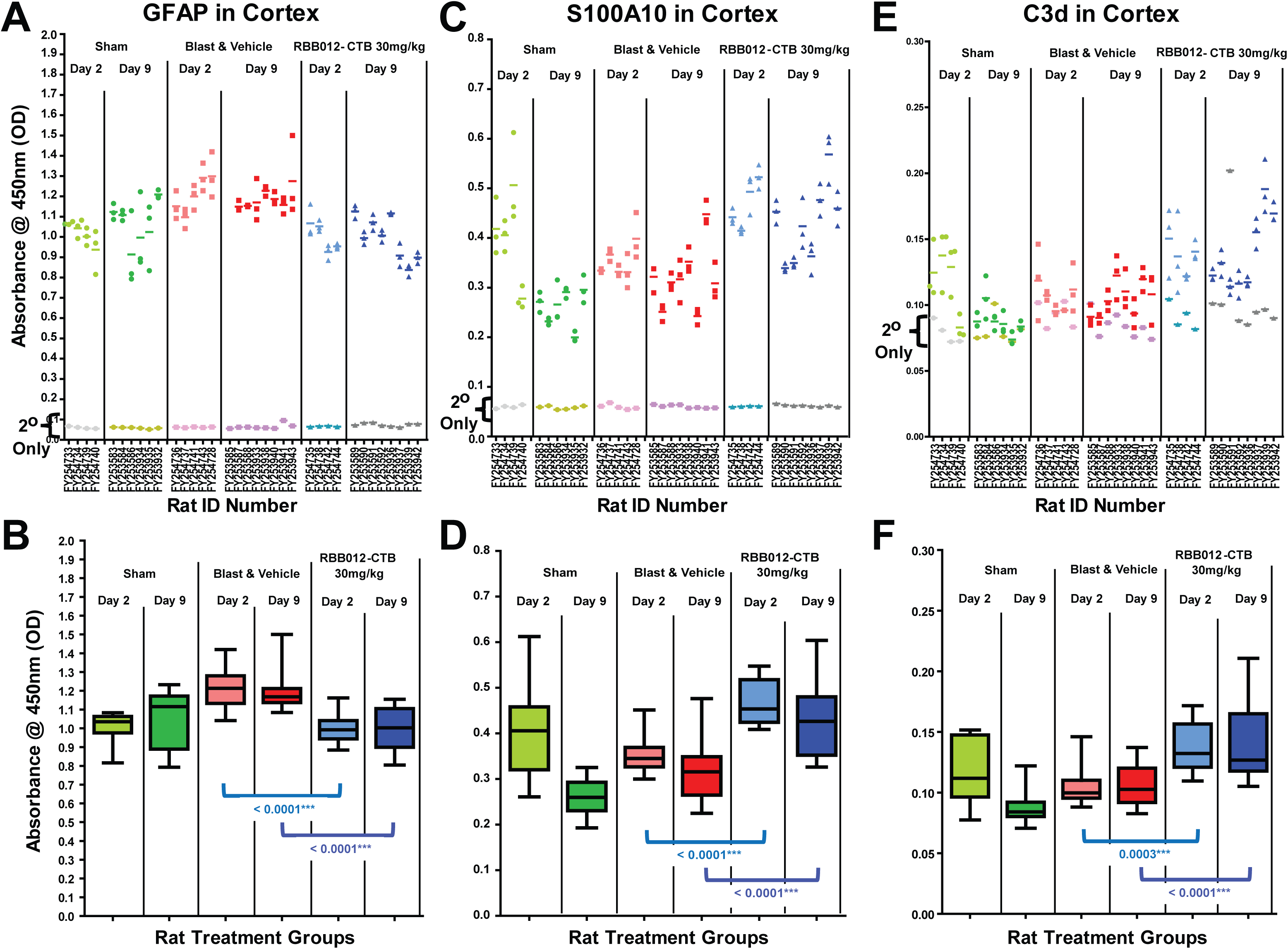
100µg of cortical tissue from the blast rats were probed, in triplicate, in the wells of an ELISA plate and probed with antibodies (**Table 1**) to characterize glial cell injury and astrogliosis. **A)** A scatter plot of the extracts probed with a rabbit polyclonal to GFAP human isotype 1 at a 1:1000 dilution, followed by a GAR-HRP secondary at a 1:2000 dilution, and TMB color development for 2.5m reveal rats treated with RBB012-CTB 15m post-blast had reduced glial injury that was statistically significant when compared to vehicle controls up to 9 days later (p<0.0001). **B)** Same results presented as box plots. **C)** A scatter plot of biomarker S100A10, shows an increase of neuroprotective astrocytes in the cortex. Extracts were probed with a mouse monoclonal at a 1:1000 dilution, followed by a GAM-HRP secondary at a 1:1000 dilution, and TMB color development for 8m. RBB012-CTB treated rats had significant increases in S100A10^+^ astrocytes compared to vehicle controls for up to 9 days later (p<0.0001). **D)** Same data presented as box plots. **E)** Neurotoxic astrocytes were detected using a goat polyclonal antibody to C3d at a 1:40 dilution and a donkey anti-goat (DAG-HRP) at 1:1000. TMB color development stopped at 15m resulted in absorbances at or near background. **F)** Even so, box plots of the same data illustrate a statistically significant increase in C3d^+^ astrocytes in those rats given RBB012-CTB up to 9 days. The same rat tissue samples incubated only with their respective secondaries (2° Only) are superimposed in each scatter plot.

### Crossing the Blood Brain Barrier

To prove drug uptake into the brain, proprietary primary sequence modifications to the HSP72 were tracked by generating several tryptic peptides that were detectable by MS as discussed elsewhere [Chan et al. *submitted*]. Briefly, three MS peptides allowed tracking of the Fv-HSP72 variants both in plasma and brain tissue. The peptide codenamed YASYL detects the 3E10 scFv subunit of RBB012 and RBB012-CTB. Peptides ITP and FGD detect sequences within the HSP72 payload allowing us to differentiate the proprietary exogenous form from its human endogenous counterpart. ITP detects HSP72 regardless of the oxidative state of the tissue it is extracted from. FGD has a methionine residue that can be oxidized. Setting the mass spectrometer to detect multiple methionine oxidation states allows evaluation of the relative oxidation exposure of the drug.

PK studies tracking both the 3E10 and HSP72 subunits involved IV tail vein injections into rats either 1) anesthetized and exposed to blast (TBI) or 2) anesthetized and returned to their cage (Naïve). Both, Naïve and TBI-exposed rats were divided into 3 groups (n=5/group) with one group euthanized 1h post-drug injection, and two other groups at 4h and 12h (**Figure 1C**). The plasma clearance of RBB012-CTB in rats injected IV either 15m after anesthesia and blast exposure or given anesthesia only are plotted both as a composite of 3 peptide biomarkers (**Figure 9A**) and individually to illustrate the Lower Limit of Quantitation (LLOQ), also known as the signal to noise (S/N) threshold (**Figures 9B-D**). A detailed discussion on the LLOQ determination process for each MS peptide can be found in Chan et al. (*submitted*). LLOQs differ for each peptide given their biochemical differences. Plasma concentrations in TBI versus Naïve rats, in the 1^st^ hour for peptides ITP (p=0.034) and FGD (p=0.027), were significantly different based on a Welch’s unpaired t-test. After that, detection of the RBB012-CTB fragments was at or below the LLOQ suggesting blood clearance.

**Figure 9.**
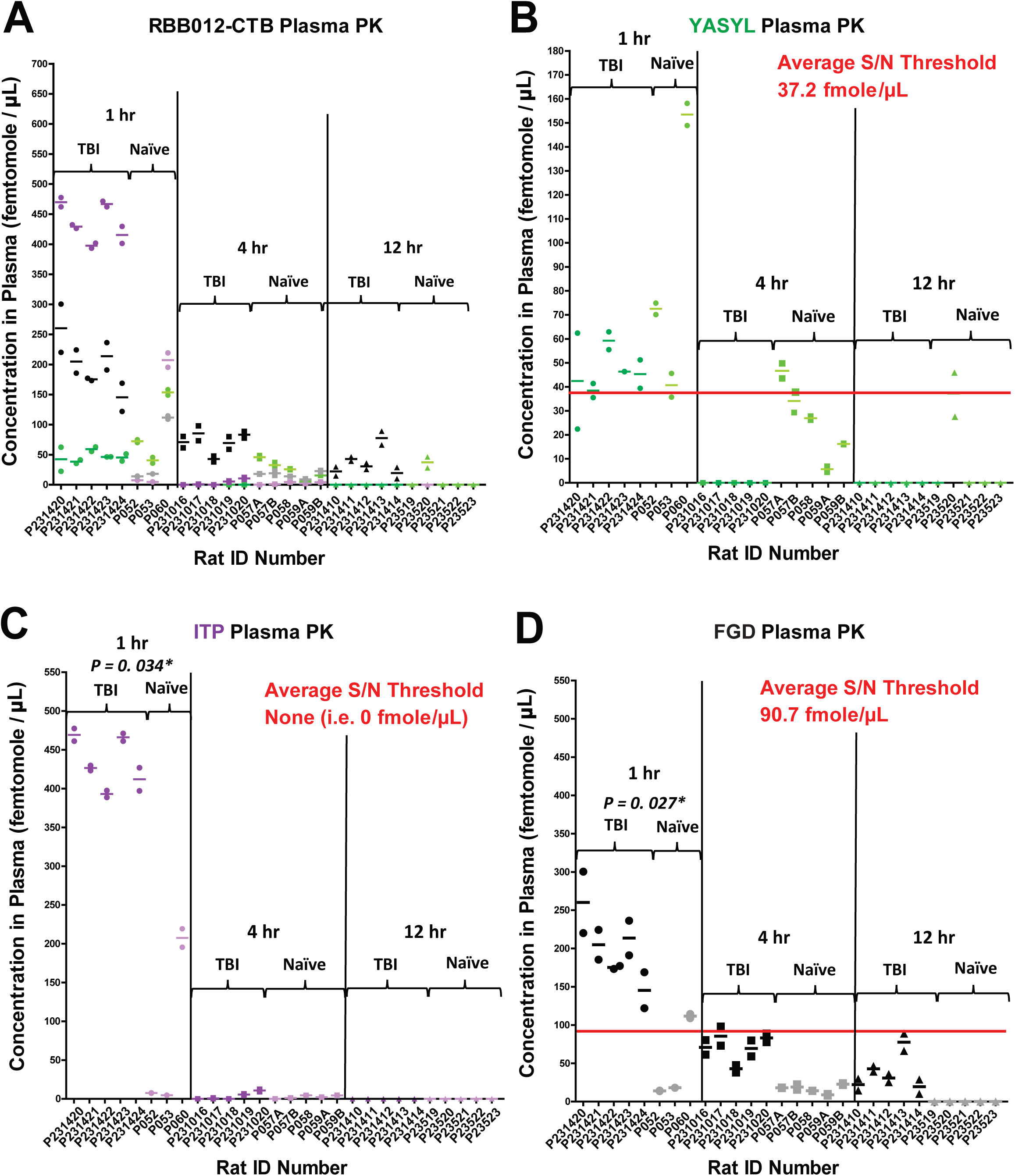
**A)** A composite graph of 3 peptide markers for RBB012-CTB helps reveal similarities in plasma concentration in the blood. Concentration of the peptide markers is plotted for each rat (whose ID numbers comprise the x-axis). Scatter plots are color coded according to the schematic in Chan et al (2026). Darker colors represent the plasma concentrations determined from rats having incurred a blast injury (TBI). Pastel colors represent the corresponding peptide concentrations from uninjured rats (“Naïve”). **B)** A scatter plot showing only the YASYL fragment helps track the variable light chain of the 3E10 subdomain comprising RBB012-CTB. Two technical repeats are plotted for each rat along with the average represented as a dashed line. An average signal:noise (S/N) threshold (i.e. the concentration most likely to comprise the LLOQ) is represented by a red line. In this case, the LLOQ is 37.2 fmole/μL. **C)** Scatter plot for the ITP peptide tracking the HSP72 domain of RBB012-CTB. **D)** Scatter plot for the FGD peptide tracking the HSP72 domain and its relative exposure to oxidation. How the LLOQ is determined for each peptide is discussed in Chan et al. (submitted).

This clearance pattern was not observed in the brain where drug was similarly retained in both TBI and Naïve rats for the first 4h post-blast. As was the case with our CCI studies, only the ITP peptide was detectable in the brain tissue above the LLOQ (**Figure 10**). Similarities in biodistribution changed significantly by 12h when the drug was neither quantifiable nor detectable in Naïve rats. We further analyzed the rate of clearance from the brain in both groups of rats before the 12h time point. To analyze peptide uptake and clearance over the first 4 hours, our statistician fit linear regression models to the ITP concentrations in TBI versus Naïve rats and used the slopes and y-intercepts for the statistical analysis. A full discussion of the method is described in Chan et al. (*submitted*). As seen in **Figure 10**, RBB012-CTB has a slightly increasing accumulation, not clearance, in Naïve rats (2.26pg/mg/h), which drops to -0.69pg/mg/h in rats exposed to a TBI. These differences are not statistically significant and should be interpreted as drug retention in the brain, rather than net accumulation; in contrast to the blood clearance seen in the same 4h period. Changes in clearance rates must be significant beyond 4h to account for the biodistribution differences at 12h.

**Figure 10.**
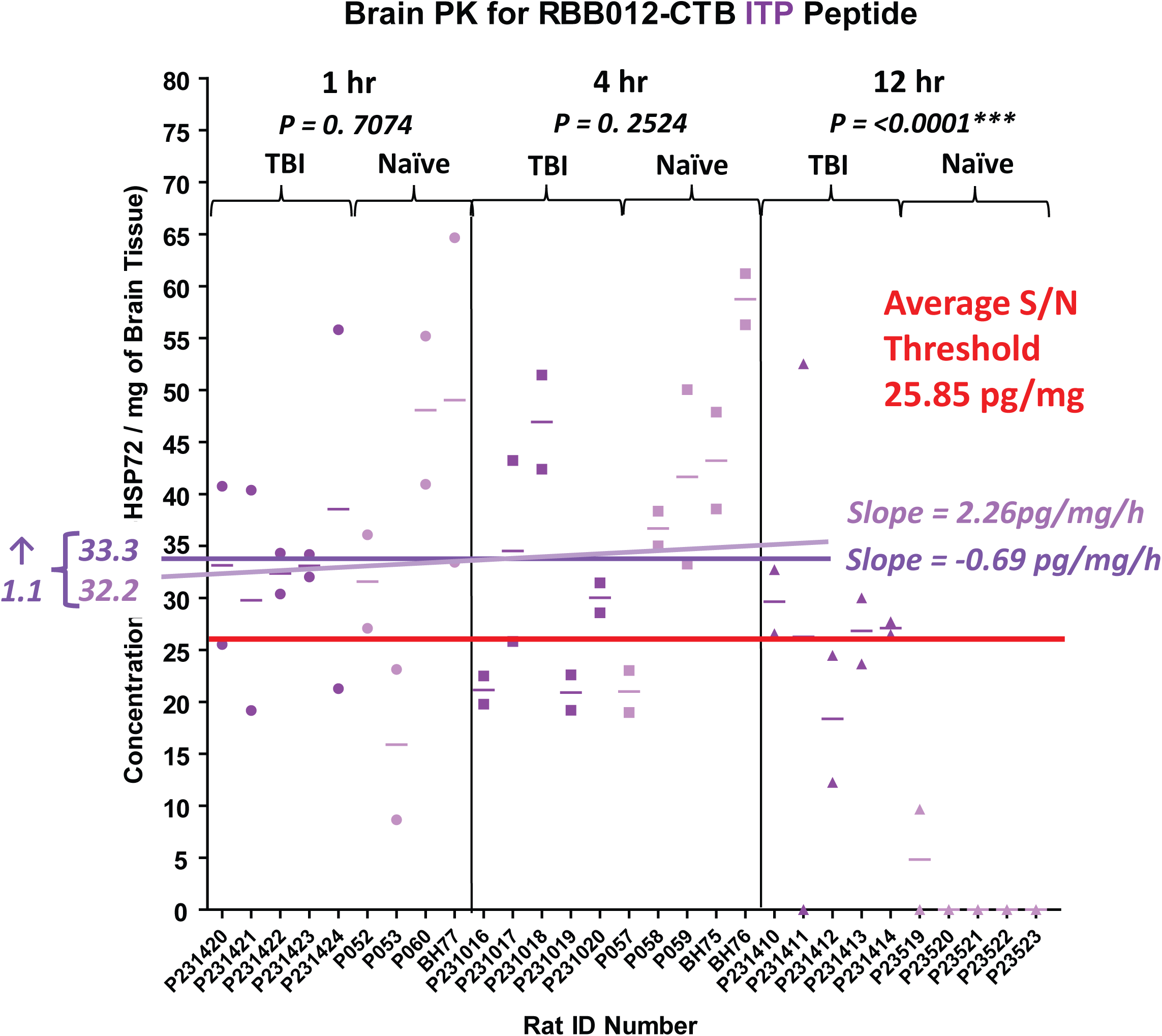
Scatter plot comparing RBB012-CTB concentration in the brain of TBI and Naïve rats. The 2 technical repeats are plotted for each rat along with the average represented as a dashed line. An average signal:noise (S/N) threshold (i.e. the concentration most likely to comprise the LLOQ) is represented by a red line. Drug concentrations reported in pg of ITP peptide per milligrams of brain tissue. Calculations to normalize the results discussed in Chan et al (2026). Normalized concentrations for TBI rats are plotted in purple, those for Naïve rats are in the lighter shade of violet. Distributions are not statistically different for TBI and Naïve rats at1hpost-IV. At 4h, when drug is already reaching unquantifiable levels in plasma, the ITP fragment of RBB012-CTB is detectable in both TBI and Naïve rats. By 12h, the ITP is undetectable in Naïve rats resulting in a significant difference compared to blast injured animals (p<0.0001). A linear regression of the ITP concentrations in Naïve rats defines a slope of 2.26pg/mg/h indicative of increased retention of the drug during the first 4h post-injection. In TBI rats, this rate drops by 2.95pg/mg/h to a clearance rate of -0.69pg/mg/h, which is not statistically significant. Projecting both slopes to the y-intercept results in starting concentrations of 32.2 and 33.3pg/mg for the Naïve and TBI rats, an insignificant difference of only 1.1pg/mg.

## Discussion

Natural HSP72 induction can take several hours in the central nervous system (CNS)^10,22^. Rapid delivery of exogenous HSP72 to brain tissue protects cells from injury^23^ and avoids the typical delays necessary for induction of the HSP72 gene. To prevent neurological damage resulting from TBI, we have tested a re-engineered HSP72 that has already shown *in vivo* efficacy in cerebral^23^ and cardiovascular^24^ infarction models. Selectivity of Fv-HSP72 *in vivo* targeting occurs because tissues undergoing significant cell injury possess a high concentration of extracellular DNA. Salvaging of the DNA by surrounding cells, through the ENT2 channel, provides 3E10 fused to HSP72 the opportunity to enter those cells and inhibit further cell death by delivering its pleiotropic payload^17^. The power of an approach targeting a wide variety of cells in an area of traumatic tissue damage cannot be overstated. Observations of HSP72 upregulation in neural, glial and *<u>vascular</u>*cells post-injury^8,10^ highlight the need to protect as many *<u>cell types</u>* as possible so as to promote rapid tissue healing and inhibit fibrosis.

Three Fv-HSP72 structural variants engineered by Rubicon were screened in a rat blast injury model at WRAIR, simulating a moderate to severe impact^19^. These studies were conducted in parallel with CCI studies at JHU [Chan et al. (*submitted*)], with tissue samples from both studies sent on dry ice to Rubicon for biomarker extraction and analysis. For each biomarker studied, all cortical tissue extracts from WRAIR and JHU were plated at the same protein concentration, ensuring comparability of results obtained from both blast and CCI injured rats.

Some biomarkers indicating treatment efficacy in the CCI model did not do so in the blast model, highlighting the fact that a direct focal impact to the cortex can generate a different pathophysiology from the more diffuse blast injury. Despite differences in the mechanism of action for CCI and blast injury models, one Fv-HSP72 variant emerged superior in treating subsequent neurodegeneration in both cases. RBB012-CTB is characterized by having a proprietary cathepsin B cleavable protein linker and showed significant efficacy in screening studies (**Figures 2-4**). As a potential clinical candidate, it was important to quickly determine if it would improve learning and memory *in vivo*; hence, use of the NOR assay^15^. A single dose given post-blast proved efficacious for up to 8d (**Figure 5**). The very same rats from this behavior study were then subjected to the same brain tissue analysis conducted for the screening studies. We focused on nerve (**Figures 6,7**) and glia (**Figure 8**), the most prevalent cells in the brain.

Tau pathology biomarkers in the brain were significantly reduced up to 9 days post-blast in rats given 30mg/kg RBB012-CTB, versus those given vehicle (**Figure 6**), when analyzed using Welch’s unpaired t-test, a more robust method than the Student’s t-test. This reduction was also seen in cervical spinal cord extracts 2d post-blast for the TBI markers (T181, T231), but was lost by 9d (**Figure 7**); perhaps not an unexpected result given the orientation of the anesthetized rats in the blast chamber, focusing the primary impact to the brain. Glial support is crucial to neural function and a rise in GFAP is considered a marker of glial cell injury^13,21^. Rats receiving drug had significantly less detectable GFAP than animals receiving vehicle controls up to 9d post-blast in the cortex (**Figure 8B**), but results were not significant in the spinal cord (data not shown). More importantly, a single RBB012-CTB IV injection may have helped direct astrogliosis in the brain toward the A2 tissue repair state up to 9d post-blast (**Figure 8D**); rather than the neurotoxic A1 state (**Figure 8F**). These results are supported by observations in the retina of rats exposed to a blast over-pressure injury (Krutik et al. *manuscript in preparation*). In light of recent observations showing astrocyte polarization toward the A2 neuroprotective state declines with aging, both in rat brain^25^ and mouse spinal cord^26^ models, our results open the door to investigating RBB012-CTB’s potential to improve neurological outcomes in adults.

Blood brain barrier disruption due to TBI has been noted starting at 30m post-injury (using MRI)^27^ and remaining permeable for up to 8h^28^. Fv-HSP72 fusion proteins are 98kD in size; thus, it was important to prove drug uptake into the brain for such a large molecule. Identification of tryptic peptides within the Fv-HSP72 carrying proprietary amino acid modifications allowed for MS tracking of RBB012-CTB. The drug cleared from the blood between 1h and 4h post-injection (**Figure 9**). Different rates of detection in the blood can be due to interactions with other molecules in the tissue matrix. For instance, lower YASYL concentrations compared to ITP and FGD may be related to random DNA binding since the peptide corresponds to a portion of the 3E10 antibody’s antigen binding region. If that is the case, determining a specific range of molecular weights to screen for using MS, given the wide diversity of DNA sizes that could be bound, would be challenging and unconvincing. While ITP and FGD are both derived from the HSP72 subunit, the lower FGD concentrations are likely due to methionine oxidation. The MS detectors had been set only to detect unoxidized FGD in this study to determine the relative oxidation conditions *in vivo*.

ITP proved most robust as it was easily detectable in both blood and brain tissues. Unlike the plasma samples, ITP was detected and quantifiable in the brain up to 12h in TBI rats (**Figure 10**), long after it was unquantifiable and largely undetectable in blood. While, lack of FGD detection may be due to total oxidation, lack of YASYL detection could be the result of 3E10, upon cell entry, being transported to the nucleus by the protease cathepsin B and then degraded^29^. A rather conservative statistical analysis focused on the 1h and 4h data shows theoretical drug uptake at Time = 0h (y-intercept) and drug clearance (slope) in Naïve and TBI rats is not significantly different. The shift in drug clearance for Naïve rat brains occurs between 4h and 12h, not unlike what was observed in CCI rats as well [Chan et al. (*submitted*)].

A question that arises is the brain accumulation of Fv-HSP72 both in TBI and Naïve rats for up to 4h. It is our opinion that this highlights the role anesthetics play in temporarily disrupting the BBB. Note, for some of the biomarker results the Sham rats had elevated absorbances comparable to the Blast Only or Blast + Vehicle controls. Naïve rats in the MS study and Sham rats in the biomarker studies were anesthetized with isoflurane even though they were not subjected to a blast. Recent research has documented BBB disruption caused by anesthetics, such as 1.4% isoflurane^30^ and propofol^31^, the latter resulting in permeability for up to 72 hours post injury. A related possibility could be the transport of Fv-HSP72 into cerebral endothelial cells through their ENT2 channels. Extracellular DNA caused by BBB disruptions may provide the conduit for access of our drug into the brain’s vasculature.

In future studies, it will be important to determine extensions of the drug administration window that deliver the same biomarker and behavioral results as an injection 15m post-blast. These proof-of-concept studies employed tissue extracts that were directly adhered to on the surfaces of a multi-well plate. Although this approach works well for solid tissue samples, it was less reproducible in plasma samples. The greater complexity and albumin-rich composition of blood hinders accurate biomarker detection on a limited plastic surface and requires a sandwich ELISA format for sensitive detection of key TBI blood biomarkers (e.g. GFAP, UCH-L1, NF-L, Tau and pTau)^13^. This format will be applied in future studies. It will also be necessary to evaluate an intranasal delivery approach, that would allow for drug administration in austere medical environments far from a clinical setting, be it in a military theater of operations or on a long space flight.

### Transparency, Rigor and Reproducibility Summary

Key aspects of the experimentation and analysis were delegated to Rubicon, WRAIR and Stanford to minimize risk of bias during the study. The Fv-HSP72s were synthesized and purified by Rubicon and their efficacy tested *in vitro* prior to transfer to WRAIR for animal testing. All rats were randomized and provided ID numbers by WRAIR. Blinded tissue samples were sent to Rubicon for the biomarker studies, while blinded samples for the mass spec studies were sent to Stanford. Prior to plotting the results, biomarker and mass spec data were unblinded by Parseghian. We performed a power analysis for the biomarker studies to determine a sample size, based upon parameter estimates of standard deviations and effect size from prior work in a focal TBI model^16^, that would allow us to see, with 80% power and a family-wise type I error rate of 0.05 (after Bonferroni correction, α=0.017 per comparison) a 25% difference in the primary endpoint between vehicle and treated groups in an experiment in which we would simultaneously compare vehicle to each of 3 drug doses. Based upon that analysis, and an assumption of a 20-25% loss due to experimental complications, we chose an N of 8 rats per group.

## Author’s Contributions

Investigation (tissue extractions & biomarker analysis): AC, KP, SS, LD, SE; Investigation (Blast model & tissue extractions): MG, CP, RJRST, GP, PA; Investigation (mass spec): KK; Conceptualization & Methodology: DH, RN, STH, KK, CLP, JRO, PA, JBL, MHP; Data Analysis (biomarker & mass spec): PA, DH, RN, KK, CLP, JBL, MHP; Data Curation: PA, MHP; Formal Analysis: PP, JRO, MHP; Resources (synthesis and purification of Fv-HSP72): RAR, AC; Project Administration: RAR, JBL, PA, MHP; Funding Acquisition: RAR, MHP; Writing & Visualization: MHP, AC; Review & Editing: PA, AC, STH, RN.

## Abbreviations

BBB: Blood Brain Barrier
CHO: Chinese hamster ovary
CCI: Controlled Cortical Impact
d day: (s)
DTT: Dithiothreitol
ENT2: equilibrative nucleoside transporter 2
Ft: foot
HSP: heat shock proteins
H: hour(s)
IV: intravenous
JHU: Johns Hopkins University
Mg: milligrams
Msec: millisecond(s)
M: minute(s)
mAb: monoclonal antibody
MRI: magnetic resonance imaging
Pg: picograms
PTSD: Post-Traumatic Stress Disorder
S: seconds
S396: Serine 396
TBS: Tris Buffered Saline
TUNEL: terminal deoxynucleotidyl transferase dUTP nick end labeling
T181: Threonine 181
T231: Threonine 231
TBI: traumatic brain injury
WRAIR: Walter Reed Army Institute of Research

## Disclaimer

The views expressed in this manuscript reflect the results of research conducted by the author(s) and do not necessarily reflect the official policy or position of the Defense Health Agency, Department of War, nor the U.S. Government. The experiments reported herein were conducted in compliance with the Animal Welfare Act and per the principles set forth in the “Guide for Care and Use of Laboratory Animals,” Institute of Laboratory Animals Resources, National Research Council, National Academy Press, 2011.

## Acknowledgements

We thank Grace Chan and Anna Su at Rubicon, Manda Saraswati at JHU and Jacquelyn Saunders at the UCLA/Vet Affairs lab for their excellent technical assistance, as well as Soren Faulkner for outstanding assistance running the hundreds of MS samples at Stanford. We also thank Sarah Fontaine, PhD and Colleen P. Lavinka, PhD of the CDMRP for helpful technical discussions. This work was partially supported through the Congressionally Directed Medical Research Programs by the Office of the Assistant Secretary of Defense for Health Affairs through the Defense Medical Research and Development Program and endorsed by the War Department **Award No. W81XWH-19-2-0027**. Opinions, interpretations, conclusions and recommendations are those of the author and are not necessarily endorsed by the War Department.

